# A hybrid geometric–feature algorithm for 2D shape similarity

**DOI:** 10.64898/2026.08.20.745909

**Authors:** Maria Evangelia Vlachou, Elizabeth Thomas, Jean Blouin

## Abstract

In this paper, we address the problem of quantifying similarity between planar 2D shapes, which is relevant to studies of internal representations in cognitive, developmental, and neurological research. We designed a set of test shapes arranged along a visually defined perceptual similarity gradient and used them to evaluate classical geometric methods for shape comparison, including Procrustes and Chamfer distance, as well as a convolutional neural network (CNN)-inspired feature-based method. Based on the limitations identified for these individual methods, we developed a hybrid Geometric-Feature Similarity (GFS) algorithm that combines geometric alignment, global contour properties, and convolutional feature-based descriptors into a unified weighted similarity score. By combining global geometric information with local structural features, the GFS algorithm more accurately reproduces human perceptual judgments of shape similarity than either geometric or feature-based methods alone. Requiring neither network training nor large labelled datasets, the proposed algorithm provides an efficient and interpretable tool for a broad range of studies involving quantitative shape comparison.

## Introduction

Two-dimensional (2D) shapes are widely used to represent spatial and structural information. They enable the characterization of object form and spatial relationships even in the absence of texture, depth, or colour cues. Quantifying shape similarity is particularly valuable in cognitive and behavioural motor research, as it enables the assessment of how spatial structures are encoded, compared, and reproduced [1]. Such measures have been particularly used in developmental studies [2], as well as in research on neurological [3] and sensorimotor disorders [4], where they provide objective tools to evaluate changes in perceptual and motor representations. Despite its apparent simplicity, measuring 2D shape similarity remains a longstanding challenge in pattern recognition due to the need to account for global geometry, local structural detail, and perceptual relevance within a unified and interpretable framework.

In many shape analysis frameworks, shapes are represented as planar curves or ordered boundary point sets, and similarity is quantified using classical geometric metrics such as Procrustes [5–7] and Chamfer distance [8,9], which measure spatial proximity based on corresponding or nearest-neighbour points, respectively. While these methods are computationally efficient and mathematically well-defined, they often exhibit limitations when applied to shapes with complex structure, non-uniform boundary sampling, or open contours. Procrustes distance relies on explicit point correspondences, making it sensitive to point ordering and sampling density, whereas Chamfer distance primarily captures global geometric proximity and may not adequately reflect local structural variations that contribute to perceived shape similarity.

Alternatively, convolutional neural networks (CNNs) have been used to learn shape representations from rasterized contour images [10]. By extracting hierarchical image features, CNN-based approaches can capture local structural characteristics such as edges, curvature, and orientation more effectively than purely geometric methods. However, they remain sensitive to global misalignment and typically require large labelled datasets for training, making them less suitable for behavioural experiments, where the number of observations is often small.

To address these limitations, we first compare the performance of classical geometric methods and a CNN-inspired feature-based approach in reproducing perceptual shape similarity. Building on this analysis, we introduce the hybrid Geometric–Feature Similarity (GFS) framework, which integrates these geometric and feature-based approaches into a unified similarity measure. By combining global geometric information with local structural features, the proposed framework provides an interpretable and computationally efficient approach for quantifying shape similarity while avoiding the need for network training or large labelled datasets.

## Dataset and shape preprocessing

The dataset consisted of 15 manually designed planar shapes generated to represent a controlled gradient of visual similarity relative to a Reference shape (Fig. 1). Each shape was represented as an ordered set of contour points defined by its *x* and *y* coordinates. Similarity between each test shape and the Reference shape (blue outline in Fig. 1) was quantified using the different approaches described below.

**Fig 1.**
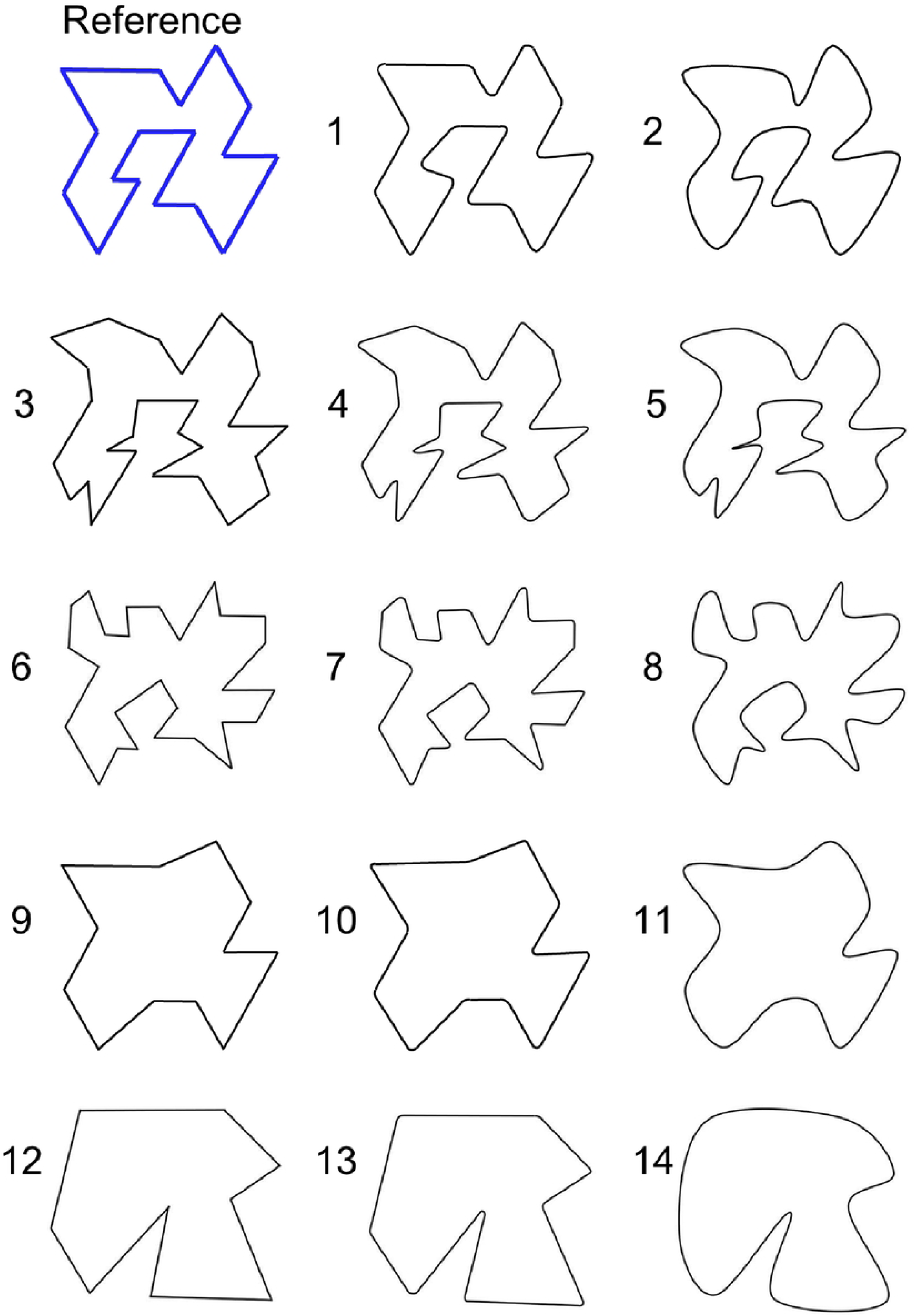
Reference shape (blue outline) and a set of test shapes ordered along a perceptual similarity gradient based on human visual judgment. Shapes are numbered from 1 (most similar) to 14 (least similar), and this numbering is used consistently throughout the article.

The test shapes were designed to vary systematically along two perceptual axes: (i) curvature, progressively increasing from left to right within each row and (ii) geometric complexity, transitioning from more complex contours (Shapes 3–8) to simpler geometries (Shapes 9–14). Shapes are numbered from 1 to 14 according to decreasing similarity to the Reference shape (Shape 1 being the most similar and Shape 14 the least similar). This numbering is used consistently throughout the article.

To ensure comparability across shapes and methods, all contours were preprocessed using the same pipeline. Each contour was resampled to a fixed number of points (*N = 200*) using arc-length parameterization to obtain uniform point distribution along the boundary. Shapes were centered using the geometric median computed with an iterative Weiszfeld algorithm to remove differences in spatial position [11].

Following this common preprocessing step, method-specific normalization and alignment procedures were applied. For methods requiring explicit geometric alignment (Chamfer distance, CNN-inspired feature similarity, and GFS algorithm), contours were additionally normalized by their RMS radial distance and aligned to the Reference shape through a coarse-to-fine optimization procedure. Alignment parameters included scaling, rotation, and cyclic point-shift optimization (for closed contours), with the optimal configuration determined by minimizing the Chamfer distance between the two-point sets. Because this dataset was designed only with respect to curvature and geometric complexity, these steps ensured that similarity estimates were invariant to arbitrary differences in translation, rotation, and scale, while allowing the framework to generalize to arbitrary shape datasets, including reproduced shapes in behavioural experiments.

For Procrustes analysis, scale and rotational normalization were not applied prior to comparison, as these transformations are estimated internally by the MATLAB *procrustes* function. Instead, after resampling and centering, all possible cyclic shifts of the contour points were evaluated. For each shift, the *procrustes* function estimated the translation, rotation, and uniform scaling that minimized the Procrustes disparity between the corresponding point sets, and the cyclic shift yielding the lowest disparity was retained.

## Evaluation of existing shape similarity methods

We first evaluated the performance of individual traditional geometric and CNN-inspired shape similarity approaches. Three methods were implemented and compared: Procrustes analysis, Chamfer distance, and a CNN-inspired feature similarity approach. All methods were implemented in MATLAB R2024b.

### Procrustes analysis

Originally introduced in 1966 [5], Procrustes analysis is a classical statistical method for shape comparison that aligns two point sets by minimizing the sum of squared distances between corresponding points after translation, rotation and scaling. Given a reference shape *X* ∈ ℝ*^Nx^*^2^ and a shape to align *Y* ∈ ℝ*^Nx^*^2^, the Procrustes problem is formulated as:

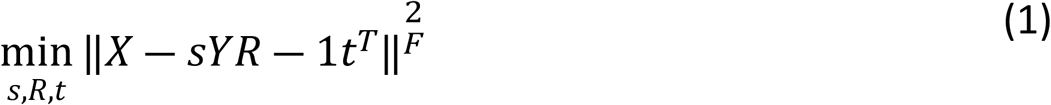

where:

- *s* ∈ ℝ is a uniform scaling factor,
- *R* ∈ ℝ^2*x*2^ is a rotation matrix (*R^T^ R* = *I*, det(*R*) = 1),
- *t* ∈ ℝ^2^ is a translation vector,
- 1 ∈ ℝ*^Nx^*^1^ is a column vector of ones to apply translation to all points,
- ∥ ⋅ ∥*_F_* denotes the Frobenius norm.

The resulting Procrustes distance quantifies the residual misalignment after the optimal translation, rotation, and scaling estimated internally by the MATLAB *procrustes* function, with lower values indicating higher similarity. To account for possible differences in point ordering, all circular shifts of the shape points were evaluated. For each shift, Procrustes alignment was performed and the configuration yielding the minimum residual error was retained.

In Fig. 2A, the best alignments of the test shapes to the Reference are shown, along with sorting by Procrustes distance.

**Fig 2.**
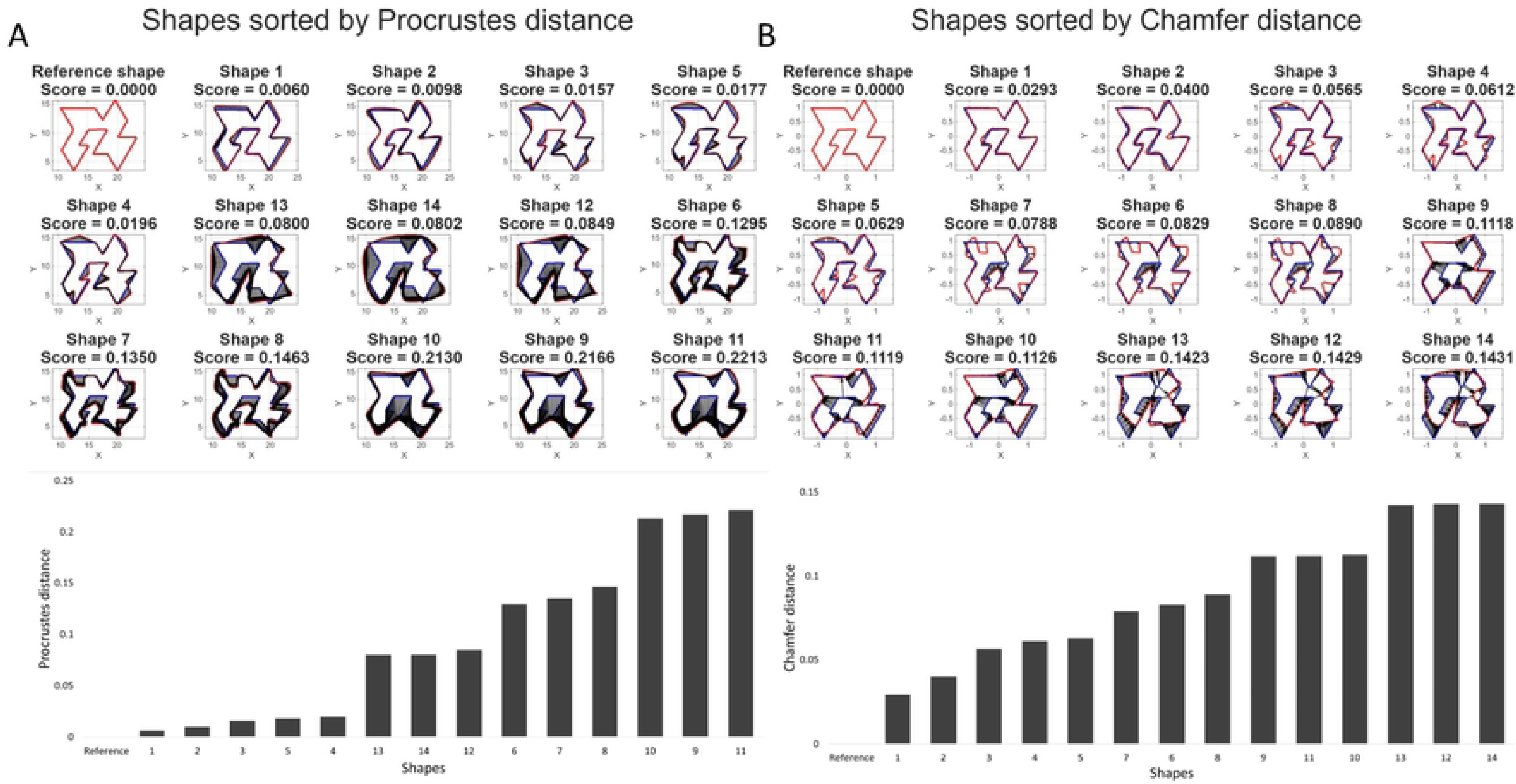
Comparison of geometric shape similarity using Procrustes analysis and Chamfer distance. **(A, Top)** Superposition of the Reference shape (blue) and aligned test shapes (red) after optimal alignment, sorted according to the Procrustes distance. Black arrows indicate point-wise error vectors. **(Bottom)** Bar plot of the Procrustes distance for each test shape relative to the Reference. Shapes are ranked in descending order of similarity score, and the shape numbering is consistent with Fig. 1. **(B)** Same representation*s* using Chamfer distance. Black arrows indicate nearest-neighbour error vectors.

### Chamfer distance

Chamfer distance is an alternative geometric measure used for comparing point sets without requiring explicit point-to-point correspondences [8,9]. It evaluates similarity by computing the average distance from each point in one set to its nearest neighbour in the other set, yielding a symmetric formulation:

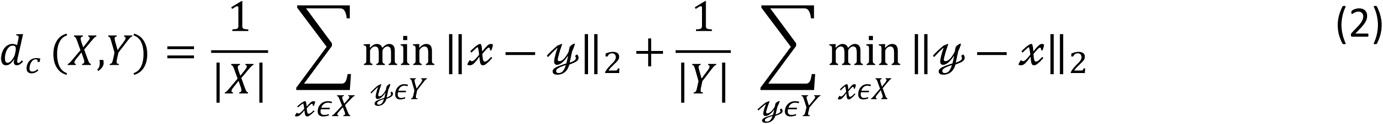

This formulation captures bidirectional nearest-neighbour proximity, providing a measure of global geometric similarity that is robust to differences in point ordering. Lower Chamfer distances indicate greater shape similarity. Unlike Procrustes analysis, which estimates optimal geometric transformations, Chamfer distance is defined as a point-set distance measure and does not inherently account for translation, rotation, or scaling. Therefore, we performed an alignment step prior to distance evaluation as described above. The aligned shapes were then ranked according to their Chamfer distance, with the corresponding optimal alignments shown in Fig. 2B.

### CNN-inspired feature similarity

CNNs have emerged as powerful tools for image analysis due to their ability to learn hierarchical feature representations directly from image data [10]. Shapes are mapped into a feature space where similarity is evaluated based on distances or correlations between learned representations. However, conventional CNN models typically require large, labelled datasets for training, which are often unavailable in cognitive and behavioural experiments where the number of observations is limited.

To address this limitation, we developed a feature-based similarity measure inspired by the early convolutional layers of CNNs [12], while using fixed handcrafted filters rather than learned kernels. Unlike point-wise geometric methods, this approach evaluates similarity in an image-based feature space by extracting multi-scale convolutional features that encode edge orientation, contour structure, and local spatial organisation. Because convolutional features are not inherently invariant to global geometric transformations, we applied the same coarse-to-fine alignment procedure used for the Chamfer distance analysis prior to feature extraction.

Following alignment, each contour was rasterized onto a fixed 64×64 grid. The contour was linearly interpolated and rendered by drawing line segments between consecutive points. The resulting image was smoothed using a Gaussian filter (σ = 2.5) to produce a continuous intensity representation. The rasterized image was normalized by its maximum intensity, preserving relative spatial structure while reducing discretization artefacts.

Five handcrafted convolutional kernels were then applied to the Reference and aligned test shapes, comprising two Sobel edge detectors (horizontal and vertical) [13], two Gaussian smoothing filters operating at different spatial scales [14], and one Laplacian-of-Gaussian (LoG) filter to capture curvature-related features [14]. For each filter *k*, the corresponding feature map was computed as:

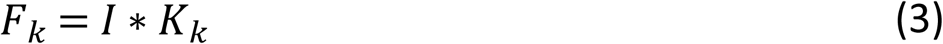

where *I* is the rasterized shape image and *K_k_* is the corresponding convolution kernel. Similarity between the Reference and test feature maps was first quantified using cosine similarity:

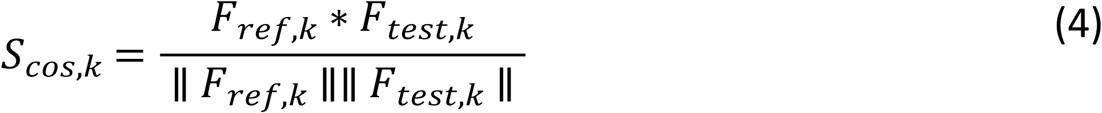

The cosine similarities were averaged across all filters to obtain a global feature similarity measure:

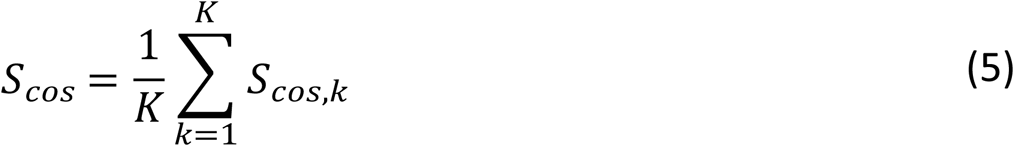

Because cosine similarity captures only the overall agreement between feature maps, an additional local spatial similarity term was introduced to preserve the spatial distribution of features. Each rasterized image was divided into a 4×4 grid, and the normalized intensity distributions within corresponding blocks were compared using the Bhattacharyya coefficient[15]:

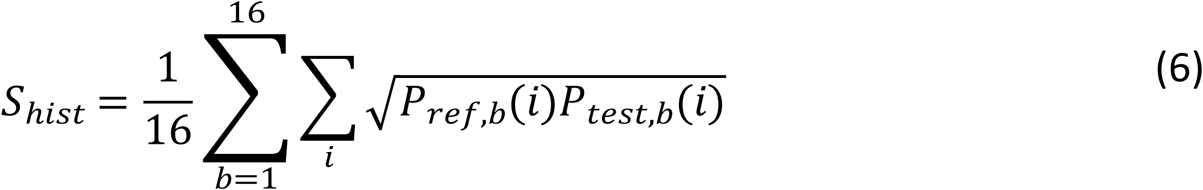

where *P_ref_*_,*b*_ and *P_test_*_,*b*_ represent the normalized intensity distributions within block *b*. The 4 × 4 partition was selected as a practical compromise between preserving coarse spatial structure and maintaining robustness to small local variations. The final feature similarity score was computed by combining the overall agreement between convolutional feature maps (*S_cos_*) with the spatial consistency of those features (*S*_ℎ*ist*_):

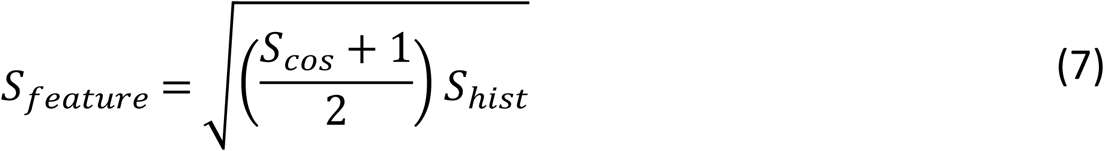

resulting in a normalized similarity value ranging from 0 to 1, where larger values indicate stronger correspondence between the Reference and test shapes.

Figure 3 illustrates a local feature similarity map derived from the aggregated absolute differences between the normalized feature maps of the Reference and aligned test shapes. For each convolutional filter, the absolute difference between the normalized feature maps was computed as:

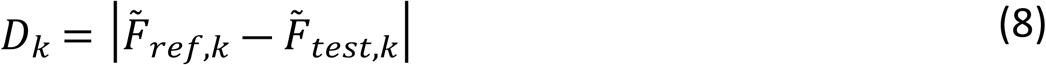

**Fig 3.**
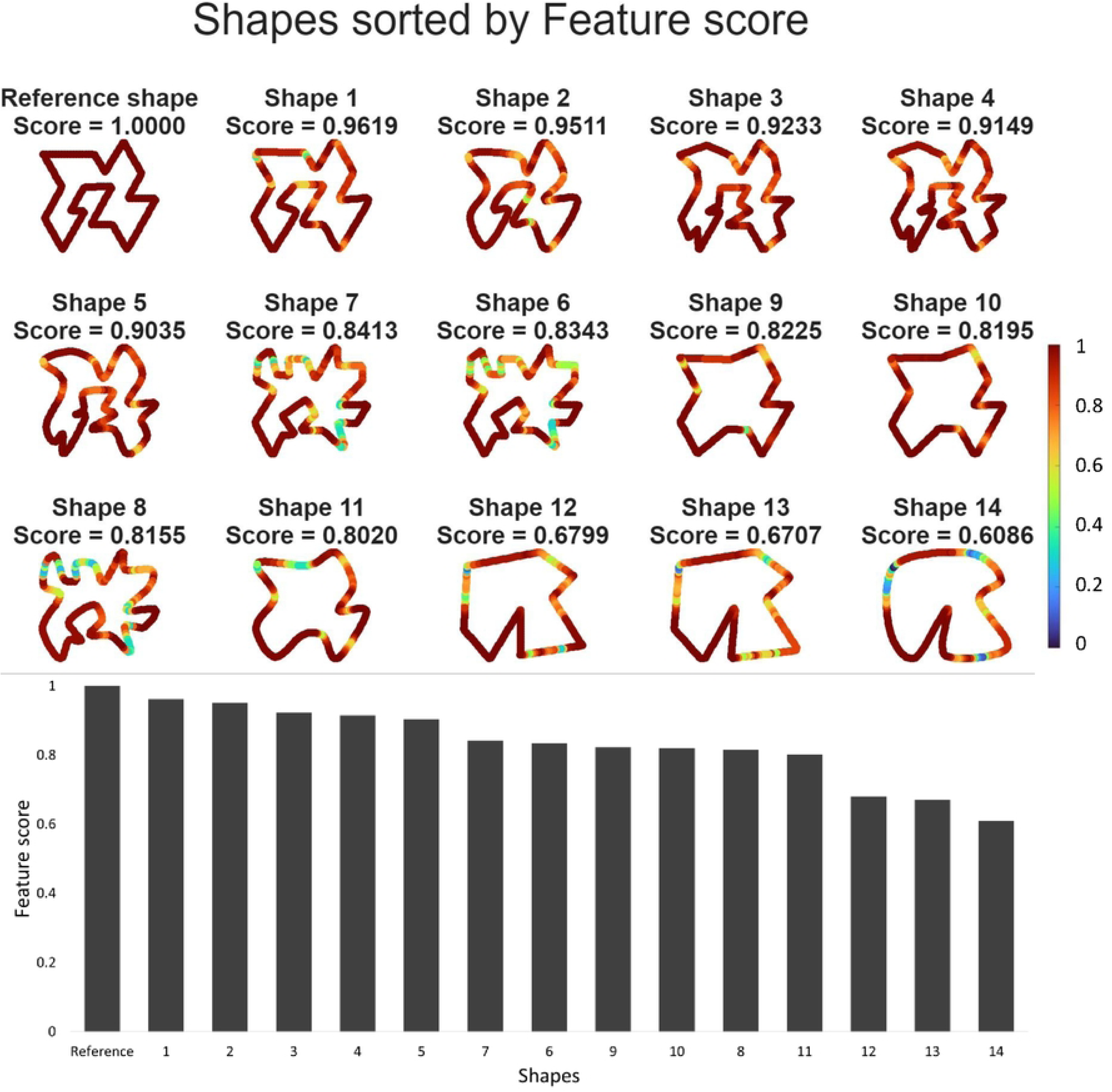
CNN-inspired feature similarity analysis. **(Top)** Heatmap visualization of local similarity between the Reference and each aligned shape. Local similarity is derived from aggregated differences between normalized convolutional feature maps and displayed using a common global colour scale, where warmer colours (red) indicate higher local similarity and cooler colours (blue) indicate lower local similarity. **(Bottom)** Bar plot of the Feature score for each test shape relative to the Reference. Shapes are ranked in descending order of similarity score, and the shape numbering is consistent with Fig. 1.

, where the feature maps were normalized using the magnitude of the corresponding Reference feature map. The final difference map was obtained by averaging across all filters,

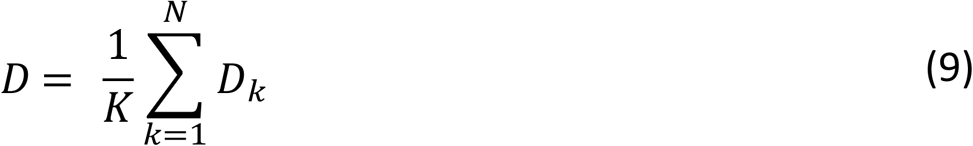

and converted into a similarity map,

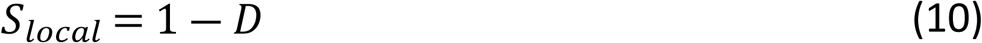

For visualization, the local similarity map was normalized using the global minimum and maximum values obtained across all analysed shapes, ensuring a common colour scale. Similarity values were then sampled at the contour coordinates of each aligned shape and displayed as a heatmap, where warmer colours (red) indicate regions of higher local agreement with the Reference shape and cooler colours (blue) indicate larger structural differences.

### Performance comparison

To quantitatively compare the performance of the three similarity methods presented above, the rankings produced by Procrustes analysis, Chamfer distance, and the CNN-inspired feature similarity method were evaluated against the perceptual ranking shown in Fig. 1. Agreement between each computational ranking and the perceptual ranking was quantified using Kendall’s rank correlation coefficient (tau), which measures the consistency of pairwise ordering between two ranked lists. Kendall’s tau ranges from −1 (complete disagreement) to 1 (perfect agreement).

Among the two geometric approaches, Procrustes analysis showed only moderate agreement with the perceptual ranking (Fig. 4A; Kendall’s tau = 0.52, p = 0.01), whereas Chamfer distance achieved substantially higher agreement (Fig. 4B; Kendall’s tau = 0.93, p < 0.001). The relatively poorer performance of Procrustes analysis likely reflects its dependence on strict point-to-point correspondences. Even after optimal alignment, local discrepancies in point correspondence can contribute disproportionately to the residual error, whereas Chamfer distance evaluates nearest-neighbour proximity and is therefore more tolerant of such local mismatches.

**Fig 4.**
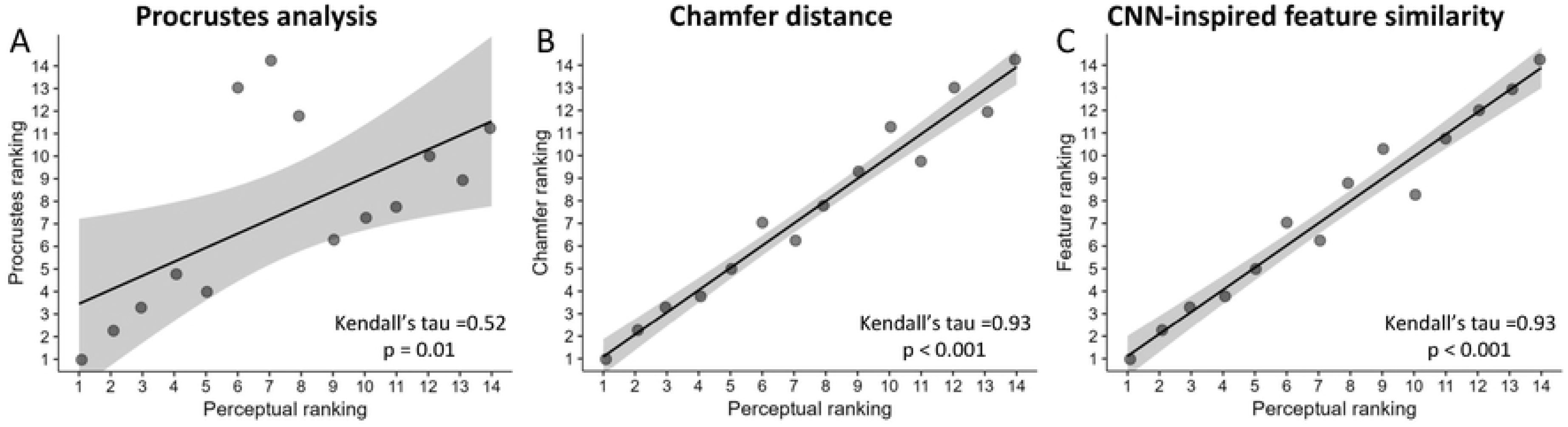
Correlation between the perceptual similarity ranking and the rankings produced by the three computational approaches. **(A)** Procrustes analysis, **(B)** Chamfer distance, and **(C)** CNN-inspired feature similarity. Each point represents one of the 14 test shapes, plotted according to its perceptual rank (x-axis) and its computational rank (y-axis). The solid line shows the linear regression fit, and the shaded region represents the standard error (SE) of the fitted regression. Kendall’s rank correlation coefficient (tau) and the corresponding p-value are reported for each method. Higher agreement with the identity relationship indicates better correspondence with the perceptual ranking.

The CNN-inspired feature similarity also showed strong agreement with the perceptual ranking (Fig. 4C; Kendall’s tau = 0.93, p < 0.001), demonstrating its ability to capture local structural characteristics.

Although Chamfer distance and CNN-inspired feature similarity achieved similarly high correspondence with the perceptual ranking, neither method was able to reproduce the human-defined ordering exactly (see bottom panel of Figs. 2 and 3). Importantly, the two approaches quantify similarity using fundamentally different representations: Chamfer distance measures geometric correspondence between aligned point sets, whereas the CNN-inspired method evaluates similarity in a convolutional feature space that emphasizes local orientation and curvature patterns. These complementary geometric and feature-based representations motivated the development of a hybrid geometric–feature similarity (GFS) algorithm, which integrates both into a unified similarity measure.

## Hybrid Geometric-Feature Similarity (GFS) algorithm

Given the limitations observed when applying each similarity measure separately, we propose a hybrid Geometric–Feature Similarity (GFS) algorithm that integrates geometric and feature-based information into a single similarity measure (see Fig. 5 for schematic pipeline).

**Fig 5.**
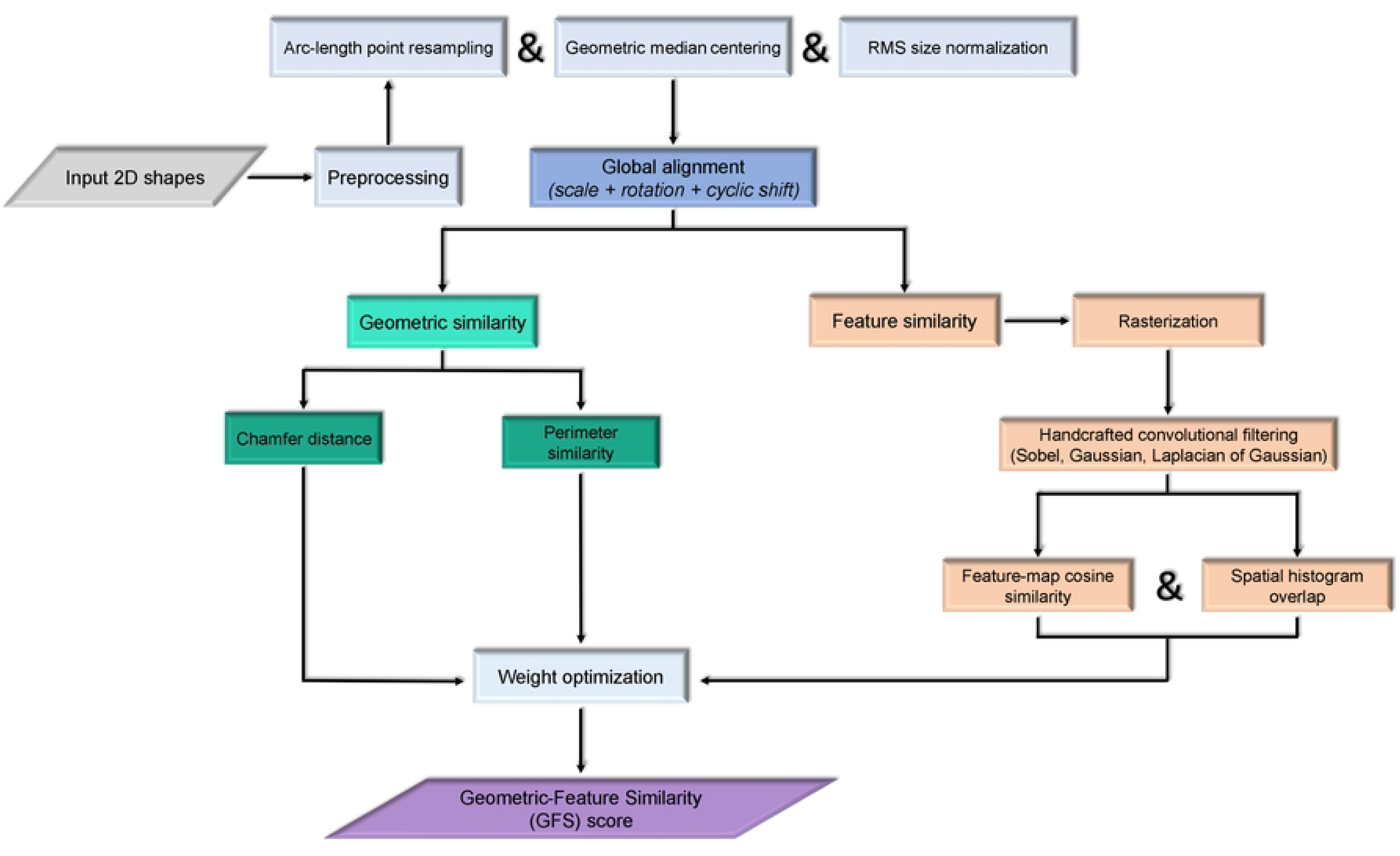
Schematic pipeline of the hybrid Geometric–Feature Similarity (GFS) algorithm. Shapes are first preprocessed by arc-length resampling, geometric median centering, and RMS normalization, followed by global alignment to the Reference through optimization of scale, rotation, and cyclic point shift. The aligned shapes are then analysed through two complementary pathways: (i) a geometric pathway, which computes Chamfer similarity and perimeter similarity, and (ii) a feature pathway, in which rasterized shapes are processed using handcrafted convolutional filters to derive a CNN-inspired feature score based on feature-map cosine similarity and spatial histogram overlap. The three similarity measures are combined into a weighted GFS score, with the weights determined by exhaustive grid-search optimization to maximize correspondence with the perceptual similarity ranking.

The GFS algorithm combines three complementary descriptors:

Geometric similarity was quantified using the Chamfer distance after optimal alignment, which outperformed Procrustes analysis in capturing global spatial correspondence between point sets (Fig. 2). To facilitate integration with the other normalized similarity measures, the Chamfer distance (*d*) was converted into a similarity score bounded between 0 and 1 using an exponential decay function:

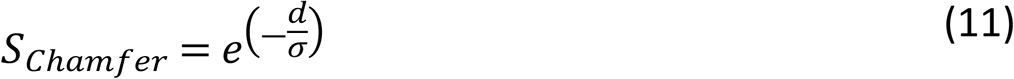

where *σ* = *s* ∗ *σ_ref_*. Here, *σ_ref_* denotes the standard deviation of the distances of the Reference contour points from the origin, and *s* is a sensitivity constant. A value of *s* = 0.4 was used, providing a scale-adaptive normalization that preserves sensitivity across different shape sizes while maintaining comparable similarity values.

To complement the Chamfer distance, we also incorporated perimeter similarity as an additional global geometric descriptor. Although two shapes may exhibit similar point-set alignment, they can differ substantially in contour length because of changes in curvature or overall geometric complexity. The perimeter similarity therefore provides a simple measure of global contour preservation that is independent of local point correspondences. It was computed as:

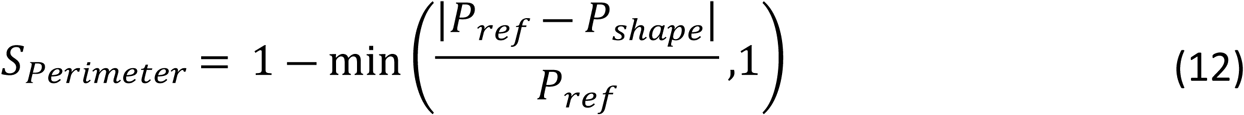

where *P_ref_* and *P_s_*_ℎ*ape*_ denote the perimeters of the Reference and aligned test shapes, respectively. This formulation yields values in the range [0,1], with 1 indicating identical contour lengths.

Feature similarity was obtained from the CNN-inspired feature extraction framework described above. This component combines the cosine similarity of convolutional feature maps with a spatial histogram overlap measure to quantify local structural similarity, including edge orientation, curvature-related contour structure, and spatial feature distribution.

The final GFS score was defined as a weighted linear combination of the three normalized similarity measures:

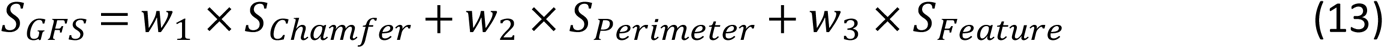

The weights were optimized automatically using an exhaustive grid search. Candidate weight combinations were sampled at intervals of 0.01 while satisfying the constraint *w*_1_ + *w*_2_ + *w*_3_ = 1. For each combination, a global GFS score was computed for every shape, and the resulting ranking was evaluated using the Normalized Discounted Cumulative Gain (NDCG). NDCG penalizes ranking errors near the top of the similarity ranking more strongly than errors occurring lower in the ranking. The optimal solution assigned weights of 0.57 to the Feature score, 0.16 to the Chamfer score, and 0.27 to the Perimeter score, achieving a perfect agreement with the perceptual ranking (NDCG = 1.00).

To assess the robustness of the optimized weights, their generalizability was evaluated using leave-one-out cross-validation (LOOCV). At each iteration, one shape was excluded, the optimal weights were re-estimated using the remaining shapes, and the left-out shape was scored using the newly derived weights. The cross-validated performance remained very high (NDCG = 0.990), with a negligible generalization gap of 0.010, indicating limited sensitivity of the optimized weighting scheme to the exclusion of individual shapes.

Figure 6 shows the aligned test shapes represented as fused heatmaps, where each point along the contour is coloured according to the local GFS score obtained by combining the normalized feature similarity map with the optimized global geometric components (Chamfer distance and perimeter similarity). Warmer colours indicate regions of higher similarity to the Reference shape, whereas cooler colours indicate lower similarity. Unlike the geometric (Procrustes, Chamfer distance) and feature-based (CNN-inspired) approaches, the hybrid GFS algorithm successfully reproduced the perceptual similarity ranking while simultaneously capturing both local structural features and global geometric organization.

**Fig 6.**
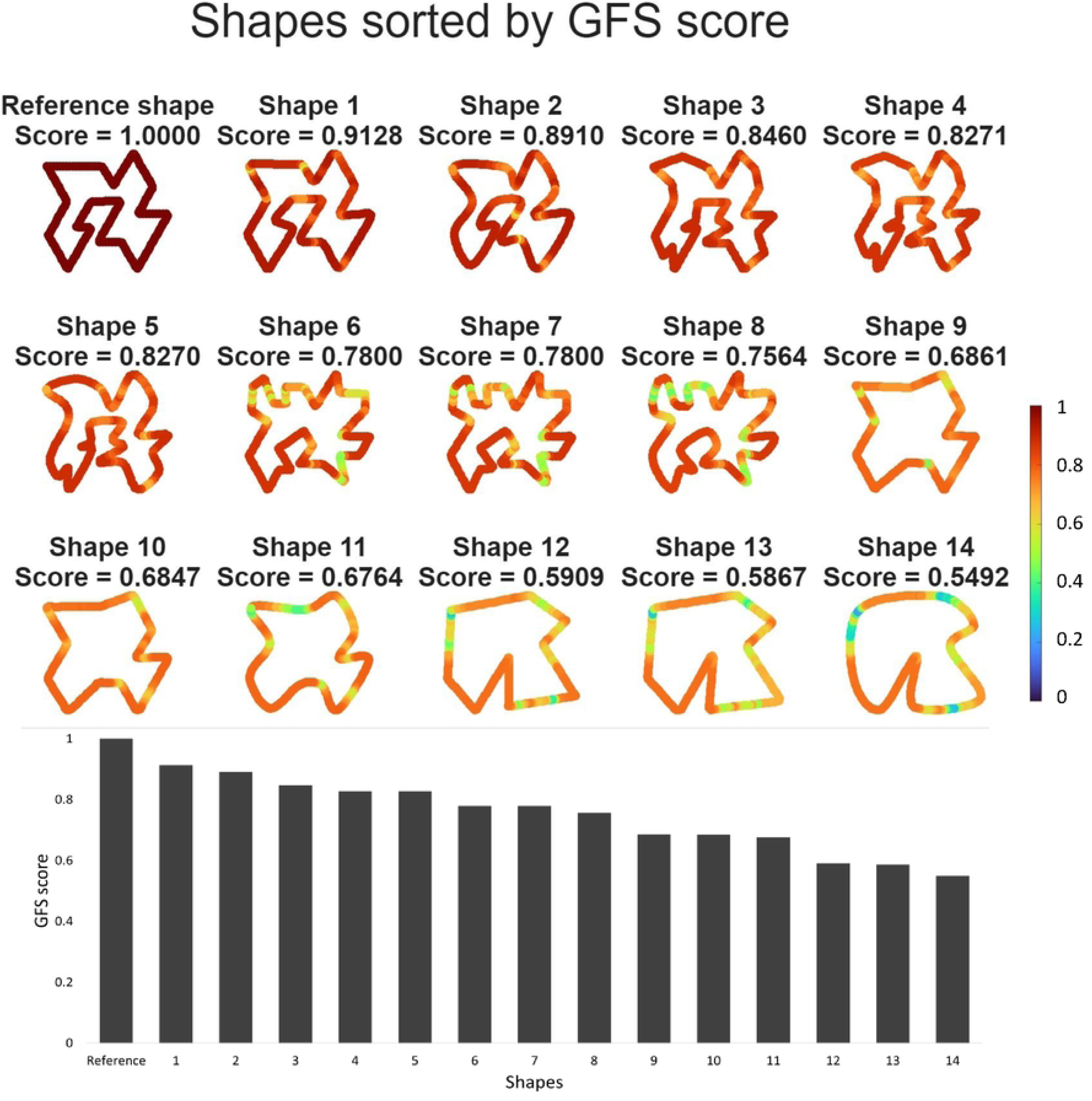
Hybrid geometric-feature similarity (GFS) algorithm. **(Top)** Heatmap visualization of local similarity between the Reference and each aligned shape. Point-wise values are obtained from the hybrid GFS score, sampled along the aligned contour, and displayed on a normalized colour scale (0–1), where warmer colours (red) indicate higher similarity to the Reference and cooler colours (blue) indicate lower similarity. **(Bottom)** Bar plot of the global GFS score for each test shape relative to the Reference. In both panels, shapes are ranked in descending order of similarity score, and the shape numbering is consistent with Fig. 1.

## Discussion

The present work introduces a hybrid Geometric–Feature Similarity (GFS) algorithm for quantifying similarity between planar 2D shapes by combining geometric and CNN-inspired feature-based descriptors. Using a set of shapes designed to follow a controlled perceptual similarity gradient, we demonstrated that although Chamfer distance and the CNN-inspired feature both showed strong agreement with the perceptual ranking, neither method alone fully reproduced the predefined ordering. By integrating global geometric correspondence, contour length, and local structural information within a single weighted similarity measure, the GFS algorithm overcame the individual limitations of these approaches and achieved perfect agreement with the perceptual ranking.

Beyond its performance, the proposed framework offers several practical advantages. The GFS algorithm is easy to implement, computationally efficient. Because it relies on handcrafted feature extraction rather than learned representations, it does not require network training or large labelled datasets. These characteristics make it particularly well suited to behavioural and perceptual experiments, where datasets are typically small, similarity is often assessed relative to a reference stimulus, and objective, reproducible similarity measures are required.

Several methodological limitations should nevertheless be acknowledged. First, the proposed implementation was designed to assess shape similarity based exclusively on contour information. Consequently, it does not incorporate interior appearance cues, such as texture, shading, or internal structures, which may also contribute to perceptual similarity. The current framework is therefore best suited to silhouette-based analyses.

Second, the GFS algorithm was evaluated using a controlled set of shapes, all of which were derived from a common Reference shape. While this design is appropriate to applications such as shape reproduction, template matching, and perceptual comparison tasks, the extent to which it generalizes to comparisons between highly dissimilar or structurally unrelated shapes remains to be established.

Finally, although the weighting scheme was determined automatically through exhaustive optimization and demonstrated excellent generalization under leave-one-out cross-validation, the optimized weights were estimated from the controlled dataset used in the present study. This is appropriate for the intended application of the framework to small-scale behavioural and perceptual experiments. Nevertheless, future work should determine whether similar weighting coefficients remain optimal across different shape classes, more complex contour geometries, and alternative perceptual paradigms.

## Code and data availability

MATLAB code and the shape dataset used in this study are available at the following GitHub repository: https://github.com/marievavl/shape-similarity-analysis.

## Authors contributions

B.J., V.M-E., and T.E. conceived the methodology. V.M-E. developed and implemented the algorithm, performed the computational analyses, and prepared the original draft of the manuscript. B.J., V.M-E., and T.E. contributed to the interpretation of the results and reviewed and edited the manuscript. All authors approved the final version of the manuscript and agree to be accountable for all aspects of the work, ensuring that questions related to the accuracy or integrity of any part of the work are appropriately investigated and resolved. All persons designated as authors qualify for authorship, and all those who qualify for authorship are listed.

